# Perceptual and Semantic Representations at Encoding Contribute to True and False Recognition of Objects

**DOI:** 10.1101/2021.03.31.437847

**Authors:** Loris Naspi, Paul Hoffman, Barry Devereux, Alexa Morcom

## Abstract

When encoding new episodic memories, visual and semantic processing are proposed to make distinct contributions to accurate memory and memory distortions. Here, we used functional magnetic resonance imaging (fMRI) and representational similarity analysis to uncover the representations that predict true and false recognition of unfamiliar objects. Two semantic models captured coarse-grained taxonomic categories and specific object features, respectively, while two perceptual models embodied low-level visual properties. Twenty-eight female and male participants encoded images of objects during fMRI scanning, and later had to discriminate studied objects from similar lures and novel objects in a recognition memory test. Both perceptual and semantic models predicted true memory. When studied objects were later identified correctly, neural patterns corresponded to low-level visual representations of these object images in the early visual cortex, lingual, and fusiform gyri. In a similar fashion, alignment of neural patterns with fine-grained semantic feature representations in the fusiform gyrus also predicted true recognition. However, emphasis on coarser taxonomic representations predicted forgetting more anteriorly in ventral anterior temporal lobe, left perirhinal cortex, and left inferior frontal gyrus. In contrast, false recognition of similar lure objects was associated with weaker visual analysis posteriorly in early visual and left occipitotemporal cortex. The results implicate multiple perceptual and semantic representations in successful memory encoding and suggest that fine-grained semantic as well as visual analysis contributes to accurate later recognition, while processing visual image detail is critical for avoiding false recognition errors.

**Significance Statement:** People are able to store detailed memories of many similar objects. We offer new insights into the encoding of these specific memories by combining fMRI with explicit models of how image properties and object knowledge are represented in the brain. When people processed fine-grained visual properties in occipital and inferior temporal cortex, they were more likely to be recognize the objects later, and less likely to falsely recognize similar objects. In contrast, while object-specific feature representations in fusiform predicted accurate memory, coarse-grained categorical representations in frontal and temporal regions predicted forgetting. The data provide the first direct tests of theoretical assumptions about encoding true and false memories, suggesting that semantic representations contribute to specific memories as well as errors.

## Introduction

Humans are able to remember objects in great detail and discriminate them in memory from others that are similar in appearance and type (Standing, 1973). To achieve this, highly specific memories must be encoded. Successful object encoding engages diverse cortical regions alongside the hippocampus (Kim, 2011). These areas intersect with networks involved in visual object processing and semantic cognition (Binder et al., 2009; Clarke and Tyler, 2014). However, little is known about the neural operations these regions support during encoding. According to fuzzy-trace theory, the specific memory traces that contribute to true recognition depend on encoding of perceptual features, while semantic gist representations promote both true and false recognition (Brainerd and Reyna, 1990). However, recent data suggest that perceptual relations between studied items and lures can also trigger false recognition (Naspi et al., 2020). Here, we used functional magnetic resonance imaging (fMRI) and representational similarity analysis (RSA) to investigate the perceptual and semantic representations engaged that allow people to recognize these same objects later among perceptually and semantically similar lures.

In line with fuzzy-trace theory, a few fMRI studies have shown stronger activation in occipito-temporal regions when people later successfully recognize specific studied objects than when they misrecognize similar lures (Garoff et al., 2005; Gonsalves et al., 2004; Okado and Stark, 2005). However, activation of similar posterior areas has also been associated with later false recognition (Garoff et al., 2005), and activation in left inferior frontal gyrus – a region typically associated with semantic processing – with later true recognition (Pidgeon and Morcom, 2016). Such results appear to challenge any simple mapping between perceptual and semantic processing and true and false recognition (see also Naspi et al., 2020). However, one cannot infer type of processing based on presence or absence of activation alone. Here, we investigated the underlying processes that give rise to such effects, using RSA to test whether patterns of neural similarity that indicate visual and semantic processing predict subsequent memory performance.

Object recognition involves visual analysis and the computation of meaning, proceeding in an informational gradient along the ventral visual pathway (Clarke and Tyler, 2015). The coarse semantic identity of an object emerges gradually from vision in posterior cortices including lingual, fusiform, parahippocampal, and inferior temporal gyri that integrate semantic features capturing taxonomic relationships (Devereux et al., 2013; Mahon et al., 2009; Tyler et al., 2013). The lingual and fusiform gyri in particular are also engaged when memories of objects are encoded (Kim, 2011). At the apex of the ventral pathway, the perirhinal cortex provides the finer-grained feature integration required to differentiate similar objects (Clarke and Tyler, 2014; Devlin and Price, 2007; Winters and Bussey, 2005), and activation here predicts later memory for specific objects (Chen et al., 2019). Other researchers ascribe this role more broadly to the anterior ventral temporal cortex, considered a semantic hub that integrates modality-specific features into transmodal conceptual representations (Lambon Ralph et al., 2017). Beyond the ventral stream, left inferolateral prefrontal regions supporting controlled, selective semantic processing are also critical for memory encoding (Gabrieli et al., 1998; Kim, 2011).

According to theory, the perceptual and semantic representations encoded in memory traces reflect how items were originally processed (Craik and Lockhart, 1972; Otten and Rugg, 2001). We therefore expected that some of these ventral pathway and inferior frontal representations would be revealed in distinct distributed activity patterns giving rise to later true and false recognition. We quantified perceptual representations in terms of low-level visual attributes of object images, and semantic representations of the objects’ concepts in terms of their coarse taxonomic category membership as well as their specific semantic features. We used these models to identify representational similarity patterns between objects at encoding using a novel approach that combined RSA and the subsequent memory paradigm in a single step. This allowed us to test where the strength of perceptual and semantic object representations predicts subsequent accurate memory and false recognition of similar lures.

## Materials and Methods

### Participants

Twenty-eight right-handed adults aged 18-35 years underwent fMRI scanning (M = 23.07 years, SD = 3.54; 18 females, 10 males). Data from a further 4 participants were excluded due to technical failures. All participants also spoke English fluently (i.e., had spoken English since the age of 5 or lived in an English-speaking country for at least 10 years) and had normal or corrected-to-normal vision. Exclusion criteria were a history of a serious systemic psychiatric, medical or a neurological condition, visual issues precluding good visibility of the task in the scanner, and standard MRI exclusion criteria (see https://osf.io/ypmdj for preregistered criteria). Participants were compensated financially. They were contacted by local advertisement and provided informed consent. The study was approved by the University of Edinburgh Psychology Research Ethics Committee (Ref. 116-1819/1). All the following procedures were pre-registered unless otherwise specified.

### Stimuli

Stimuli were pictures of objects corresponding to 491 of the 638 basic-level concepts in The Centre for Speech, Language and the Brain concept property norms (the CSLB norms; Devereux et al., 2014). These were members of 24 superordinate categories (*Appliance*, *Bird*, *Body Part*, *Clothing*, *Container*, *Drink*, *Fish*, *Flower*, *Food*, *Fruit*, *Furniture*, *Invertebrate*, *Kitchenware*, *Land Animal*, *Miscellaneous, Music*, *Sea Creature*, *Tool*, *Toy*, *Tree*, *Vegetable*, *Vehicle*, *Water Vehicle*, *Weapon*), and 238 were living things and 253 non-living things. We sourced two images for each basic-level concept. Of the 982 images, 180 were a subset of the images used by Clarke and Tyler (2014), 180 were compiled from the Bank of Standardized Stimuli (BOSS; Brodeur et al., 2014) and the remaining 622 were taken from the Internet. Each study list included single exemplar images of either 328 or 327 concepts. Of these, half were subsequently tested as old and half were subsequently tested test as lures. Each test list consisted of 491 items: 164 (or 163) studied images, 164 (or 163) similar lures (i.e., images of different exemplars of studied basic-level concepts), and 163 (or 164) novel items (i.e., images of basic-level concepts that had not been studied). Three filler trials prefaced the test phase. For each participant, living and non-living concepts were randomly allocated to the conditions with equal probability, i.e., to be studied/lure or novel items. Each study and test list was presented in a unique random trial order.

### Procedure

The experiment comprised a scanned encoding phase followed by a recognition test phase outside the scanner. Stimuli were presented using MATLAB 2019b (The MathWorks Inc., 2019) and PsychToolbox (Version 2.0.14; Kleiner et al., 2007). In the scanner, stimuli were viewed on a back-projection screen via a mirror attached to the head coil. Earplugs were employed to reduce scanner noise, and head motion was minimized using foam pads. During the study phase participants viewed one image at a time, and they were asked to judge whether the name of each object started with a consonant or with a vowel, responding with either index finger via handheld fiber-optic response triggers. By requiring participants to retrieve the object names, we ensured that they processed the stimuli at both visual and semantic levels. Participants were not informed of a later memory test. Images were presented centrally against a white background for 500 ms. This was followed by a black fixation cross with duration sampled from integer values of 2 to 10 s with a flat distribution, and then a red fixation cross of 500 ms prior to the next trial, for a stimulus onset asynchrony (SOA) of 3-11 s (M = 6). At test, participants viewed one image at a time for 3 s followed by a black fixation cross for 500 ms, and they judged each picture as “old” or “new” indicating at the same time whether this judgment was accompanied by high or low confidence using one of 4 responses on a computer keyboard. Mappings of responses to hands were counterbalanced at both encoding and retrieval.

### fMRI acquisition

Images were acquired with a Siemens Magnetom Skyra 3T scanner at the Queen’s Medical Research Centre (QMRI) at the Royal Infirmary of Edinburgh. T2*-weighted functional images were collected by acquiring multiple echo-time sequences for each echo-planar functional volume (repetition time (TR) = 1700 ms, echo time (TE) = 13 ms (echo-1), 31 (echo-2) ms, and 49 ms (echo-3)). Functional data were collected over 4 scanner runs of 360 volumes, each containing 46 slices (interleaved acquisition; 80 × 80 matrix; 3 mm × 3 mm × 3 mm, flip angle = 73°). Each functional session lasted ∼ 10 min. Before functional scanning, high-resolution T1-weighted structural images were collected with TR = 2620 ms, TE = 4.9 ms, a 24-cm field of view (FOV), and a slice thickness of 0.8-mm. Two field map magnitude images (TE = 4.92 ms and 7.38 ms) and a phase difference image were collected after the 2^nd^ functional run. At the end, T2-weighted structural images were also obtained (TR= 6200 ms and TE = 120 ms).

### Image preprocessing

Except where stated, image processing followed procedures preregistered at https://osf.io/ypmdj and was conducted in SPM 12 (v7487) in MATLAB 2019b. The raw fMRI time series were first checked to detect artefact volumes that were associated with high motion or were statistical outliers (e.g. due to scanner spikes). We checked head motion per run using an initial realignment step, classifying volumes as artefacts if their absolute motion was > 3 mm or 3 deg, or between-scan relative motion > 2 mm or 2 deg. Outlier scans were then defined as those with normalized mean or standard deviation (of absolute values or differences between scans) > 7 SD from the mean for the run. Volumes identified as containing artefacts were replaced with the mean of the neighboring non-outlier volumes, or removed if at the end of a run. If more than half of the scans in a run had artefacts, that run was discarded. Artefacts were also modeled as confound regressors in the first level design matrices. Next, BOLD images acquired at different echo times were realigned and slice time corrected using SPM12 defaults. The resulting images were then resliced to the space of the first volume of the first echo-1 BOLD time series. A brain mask was computed based on preprocessed echo-1 BOLD images using Nilearn 0.5.2 and combined with a grey-and-white matter mask in functional space for better coverage of anterior and ventral temporal lobes (Abraham et al., 2014). The three echo time series were then fed into the Tedana workflow (Kundu et al., 2017), run inside the previously created brain mask. This workflow decomposed the time series into components and classified each component as BOLD signal or noise. The three echo series were optimally combined and noise components discarded from the data. The resulting time series were unwarped to correct for inhomogeneities in the scanner’s magnetic field: the voxel displacement map calculated from the field maps was coregistered to the first echo-1 image from the first run, and applied to the combined time series for each run. The preprocessed BOLD time series corresponding to the optimal denoised combination of echoes outputted by the Tedana workflow were then used for RSA analysis, where we used unsmoothed functional images in native space to keep the finer-grained structure of activity. For univariate analysis, the preprocessed BOLD time series were also spatially normalized to MNI space using SPM’s non-linear registration tool, DARTEL; spatially normalized images were then smoothed with an 8 mm isotropic full-width half maximum Gaussian kernel.

### Experimental design and statistical analysis

#### Sample size

The sample size was determined using effect sizes from two previous studies. Staresina et al. (2012) reported a large encoding-retrieval RSA similarity effect (d = 0.87). However, subsequent memory effects are typically more subtle, for example d = 0.57 for an activation measure (Morcom et al., 2003). We calculated that, with N = 28, we would have .8 power to detect d = 0.55 for a one sample *t*-test at alpha = 0.05 (G*Power 3.1.9.2).

#### Behavioral analysis

To assess whether differences in task engagement during memory encoding predicted later memory, we modelled the effects of encoding task accuracy (0, 1) on subsequent memory outcomes using two separate generalized linear mixed effect models (GLMM) for studied items tested as old (subsequent hits and misses as predictors), and for studied items tested as lures (subsequent false alarms and correct rejection as predictors). Similarly, to assess any differences in study phase reaction times (RTs) according to subsequent memory status, we used two further linear mixed effect models (LMM). At test, to evaluate the effects on memory of perceptual and semantic similarity between objects, we also applied a generalized linear mixed model following the methods of Naspi et al. (2020). This had dependent measures of response at test (“old” or “new”) and confusability predictors calculated for each image and concept. C1 visual and color confusability were defined as the similarity value of an image with its most similar picture (i.e., the nearest neighbor) from Pearson correlation and earth’s mover distance metrics, respectively. Concept confusability was calculated by a weighted sum of the cosine similarities between objects in which each weight was the between-concept similarity itself, i.e., the sum of squared similarities (see Naspi et al., (2020)). All the analyses described above were carried out data with the lme4 package (Version 1.1-23) in R (Version 4.0.0). Models included random intercepts to account for variation over items and participants.

### Multivariate fMRI analysis

#### Overview

The goal of our study was to investigate how perceptual and semantic representations processed at encoding predict successful and unsuccessful mnemonic discrimination. To test this, we used RSA to assess whether the fit of perceptual and semantic representational models to activity patterns at encoding predicted subsequent memory. In two main sets of analyses we examined representations predicting later true recognition of studied items, and representations predicting false recognition of similar lures. We implemented a novel approach that models the interaction of representation similarity with subsequent memory in a single step. Each memory encoding model contrasts the strength of visual and semantic representations of items later remembered versus forgotten (or falsely recognized versus correctly rejected) within the same representational dissimilarity matrix (RDM). In a third set of analyses we also aimed to replicate Clarke and Tyler (2014) key findings regarding perceptual and semantic representations irrespective of memory. All RSA analyses were performed separately for each participant on trial-specific parameter estimates from a general linear model (GLM). We then followed three standard steps: 1) For each theoretical perceptual and semantic model, we created model RDMs embodying the predicted pairwise dissimilarity over items; 2) For each ROI (or searchlight sphere), we created fMRI data RDMs embodying the actual dissimilarity of multivoxel activity patterns over items; 3) We determined the fits between the model RDMs and the fMRI data RDM for each ROI (or searchlight sphere). The implementation of these steps is outlined in the following sections.

#### RSA first level general linear model

Statistical analysis of fMRI data was performed in SPM12 using the first-level GLM and a Least-Squares-All (LSA) method (Mumford et al., 2012). For each participant, the design matrix included one regressor for each trial of interest, for a total of 327 or 328 regressors (depending on counterbalancing), computed by convolving the 0.5 s duration stimulus function with a canonical hemodynamic response function (HRF). For each run, we also included twelve motion regressors comprising the three translations and three rotations estimated during spatial realignment, and their scan-to-scan differences, as well as individual scan regressors for any excluded scans, and session constants for each of the 4 scanner runs. The model was fit to native space pre-processed functional images using Variational Bayes estimation with an AR(3) autocorrelation model (Penny et al., 2005). A high-pass filter with a cutoff of 128 s was applied and data were scaled to a grand mean of 100 across all voxels and scans within sessions. Rather than using the default SPM whole-brain mask (which requires a voxel intensity of 0.8 of the global mean and can lead to exclusion of ventral anterior temporal lobe voxels), we set the implicit mask threshold to 0 and instead included only voxels which had at least a 0.2 probability of being in grey or white matter, as indicated by the tissue segmentation of the participant’s T1 scan.

#### Regions of interest

All regions of interests (ROIs) are shown in Figure 1. We defined six ROIs including areas spanning the ventral visual stream, which have been implicated in visual and semantic feature-based object recognition processes (Clarke and Tyler, 2014; Clarke and Tyler, 2015). We also included the left inferior frontal gyrus, strongly implicated in semantic contributions to episodic encoding (Kim, 2011), and bilateral anterior ventral temporal cortex, which is implicated in semantic representation (Lambon Ralph et al., 2017) and is hypothesized to contribute to false memory encoding, albeit mainly in associative false memory tasks (Chadwick et al., 2016; Zhu et al., 2019). Except where explicitly stated, ROIs were bilateral and defined in MNI space using the Harvard-Oxford structural atlas: 1) the early visual cortex (EVC; BA17/18) ROI was defined using the Julich probabilistic cytoarchitectonic maps (Amunts et al., 2000) from the SPM Anatomy toolbox (Eickhoff et al., 2005); 2) the posterior ventral temporal cortex (pVTC) ROI consisted of the inferior temporal gyrus (occipito-temporal division; ITG), fusiform gyrus (FG), lingual gyrus (LG), and parahippocampal cortex (posterior division; PHC); 3) the perirhinal cortex (PrC) ROI was defined using the probabilistic perirhinal map including voxels with a > 10% probability to be in that region (Devlin and Price, 2007; Holdstock et al., 2009); 4) the anterior ventral temporal cortex (aVTC) ROI included voxels with >30% probability of being in the anterior division of the inferior temporal gyrus and >30% probability of being in the anterior division of the fusiform gyrus; 5) the left inferior frontal gyrus (LIFG; BA44/45) consisted of the pars triangularis and pars opercularis. Lastly, we used univariate analysis as a preregistered method to define additional ROIs for RSA around any regions not already in the analysis that showed significant subsequent memory effects. Based on this analysis, we also included 6) the left inferior temporal gyrus (occipito-temporal division; LITG) as defined using the Harvard-Oxford atlas (see Results, Univariate fMRI analysis). The LITG has been previously implicated in true and false memory encoding (Dennis et al., 2007; Kim and Cabeza, 2007). The ROIs in Figure 1 are mapped on a pial representation of cortex using the Connectome Workbench (https://www.humanconnectome.org/software/connectome-workbench).

**Figure 1.**
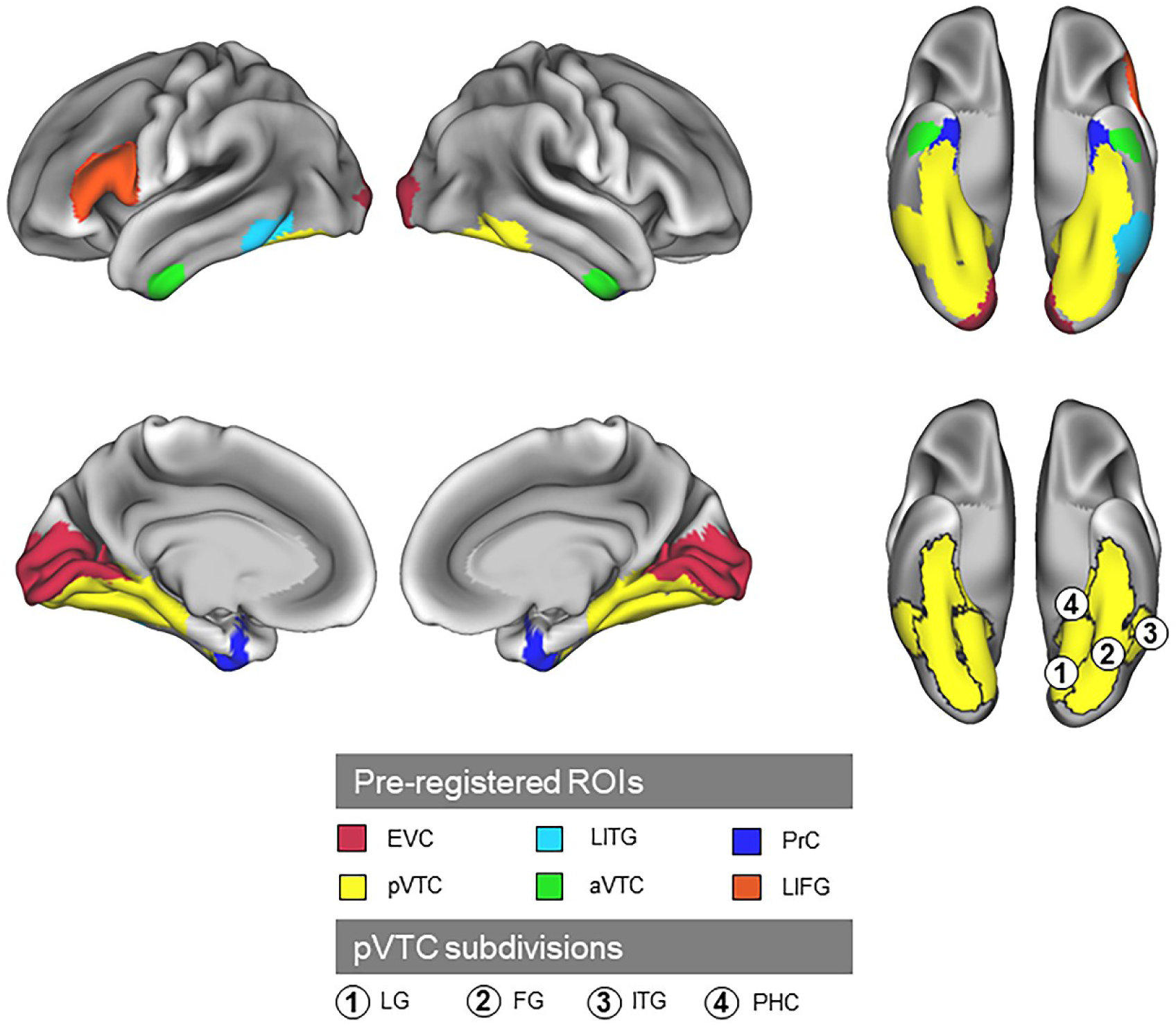
Binary ROIs overlaid on a pial cortical surface based on the normalized structural image averaged over participants. Colored ROIs represent regions known to be important in episodic encoding and in visual or semantic cognition. Circled numbers specify different subregions within pVTC (see Region of Interest for details).

#### RSA region of interest analysis

Model RDMs. We created four theoretical RDMs using low-level visual, color, binary-categorical, and specific object semantic feature measures. Figure 2 illustrates the multidimensional scale (MDS) plots for the perceptual and semantic relations expressed by these models, and Figure 3 shows the model RDMs. Memory encoding RDMs are displayed in Figure 3A and 3B, and overall RDMs irrespective of memory in Figure 3C.

**Figure 2.**
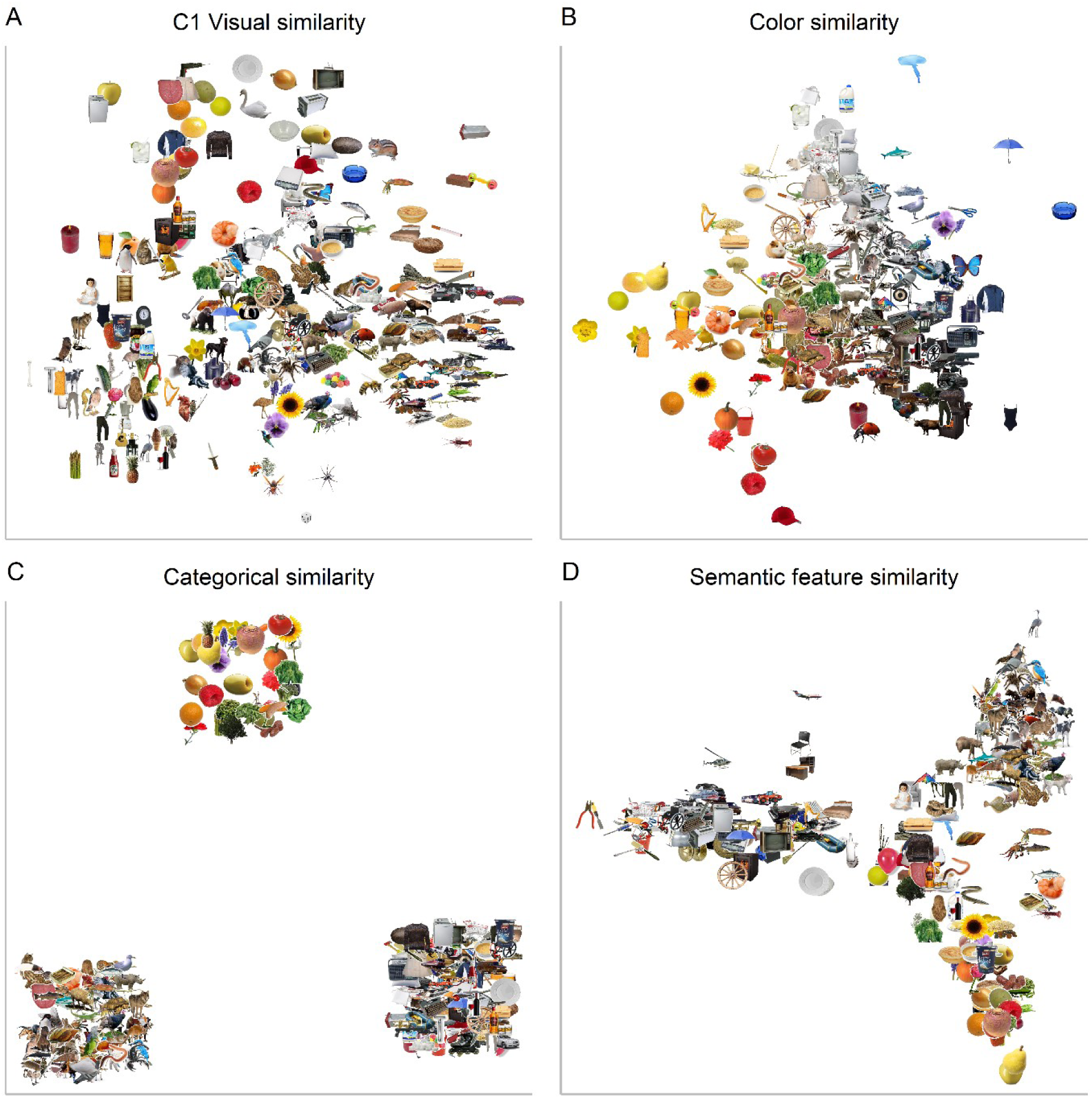
MDS plots for perceptual and semantic similarities for the four models. Pair-wise similarities were calculated to create representational dissimilarity matrices (RDMs). **A**, C1 visual similarity codes for a combination of orientation and shape (e.g., round objects towards the top, horizontal shapes on the right, vertical shapes at the bottom). **B**, Color similarity represents color saturation and size information (i.e., from bright on the left to dark at the bottom, and white towards the top). **C**, Binary categorical semantic similarity codes for domain-level representations distinguishing animals, plants and nonbiological objects (bottom-left, top, bottom-right, respectively). **D**, Semantic feature similarity codes for finer-grained distinctions based on features of each concept (e.g., differences within living things at the bottom, non-living things on the left, and many categories of animal on the top-right). The objects shown are taken from a single subject at encoding.

**Figure 3.**
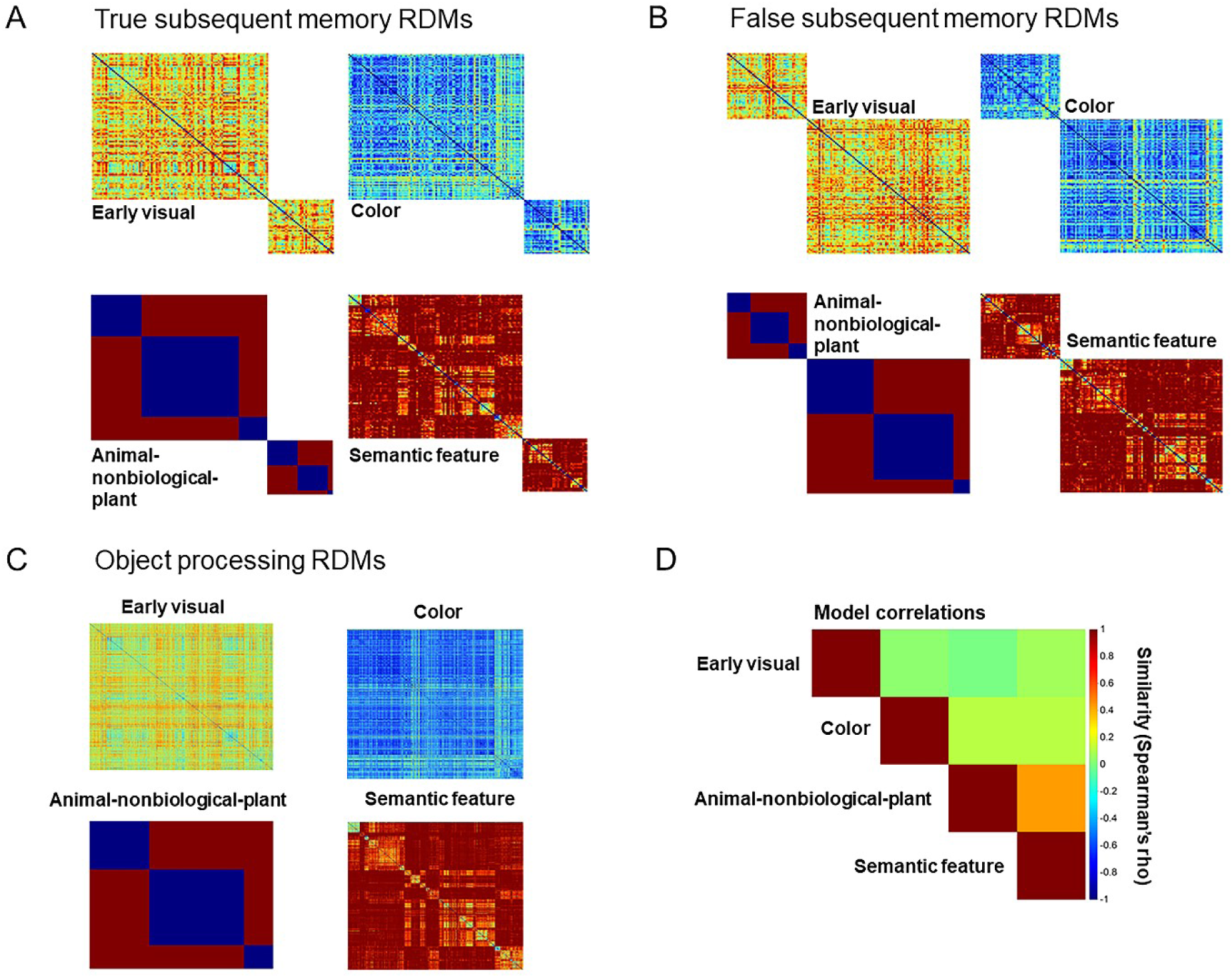
Representational dissimilarity matrices. **A**, Dissimilarity predictions of the four true subsequent memory models which included items that were later tested as old, coding subsequent hits positively (upper-left quadrants) and subsequent misses negatively (bottom-right quadrants). **B**, Dissimilarity predictions of the four false subsequent memory models which included items that were later tested as lures, coding subsequent false alarms positively (upper-left quadrants) and subsequent correct rejections negatively (bottom-right quadrants). **C**, Dissimilarity models of object processing including all the items. **D**, Similarity between theoretical models. The specific models are unique for each participant. For visualization purposes, similarity values within true and false subsequent memory RDMs have not been scaled.

1) The *early visual RDM* was derived from the HMax computational model of vision (Riesenhuber and Poggio, 1999; Serre et al., 2007) and captured the low-level (V1) visual attributes of each picture in the C1 layer. Pairwise dissimilarity values were computed as 1 - Pearson’s correlations between response vectors for gray-scale versions of each image.

2) The *color RDM* was calculated using the color distance package (Version 1.1.0; Weller and Westneat, 2019) in R. After converting the RGB channels into CIELab space we calculated the earth mover’s distance between each pair of images (Rubner et al., 2000). We then normalized the distance so that the dissimilarity values ranged from 0 (lowest) to 1 (highest).

3) The *animal-nonbiological-plant RDM* combined the 24 object categories together according to 3 domains: animal, nonbiological, and plants (Clarke and Tyler, 2014). Pairwise dissimilarity values in this RDM were either 0 (same domain) or 1 (different domain).

4) Construction of the *semantic feature RDM* followed Clarke and Tyler (2014), but used updated property norms (Devereux et al., 2014). We first computed pairwise feature similarity between concepts from a semantic feature matrix in which each concept is represented by a binary vector indicating whether a given feature is associated with the concept or not. Pairwise dissimilarity between concepts was computed as 1 – S where S is equal to the cosine angle between feature vectors. This RDM captures both categorical similarity between objects (as objects from similar categories have similar features) and within-category object individuation (as objects are composed of a unique set of features).

For the analyses of memory encoding, model RDMs were split into two, giving one RDM for each subsequent memory analysis. The true subsequent memory RDMs included only items that were subsequently tested as old; these were coded as subsequent hits or subsequent misses (Fig. 3A). The false subsequent memory RDMs included only items that were subsequently tested as lures; these were coded as subsequent false alarms or subsequent correct rejections (Fig. 3B). For true subsequent memory, we computed dissimilarity between all pairs of subsequently remembered items, and all pairs of subsequently forgotten items, omitting pairings of subsequently remembered and subsequently forgotten items. Then, to assess how dissimilarity depended on subsequent memory we weighted the model RDMs so that the sum of the cells corresponding to remembered items equaled 1 and the sum of the cells corresponding to forgotten items equaled -1, so the dissimilarity values for all included trials summed to 0 (i.e., subsequent hits – subsequent misses). Thus, positive correlations of the model RDMs with the fMRI data RDMs indicate that the representations are aligned more strongly with neural patterns for items that are later remembered than forgotten. Conversely, negative correlations indicate greater alignment for items that are later forgotten than remembered items. For false subsequent memory, we followed the same procedure, but subsequent false alarms were substituted for subsequent hits, and subsequent correct rejections for subsequent misses. Analyses were implemented using custom MATLAB 2019b (The MathWorks Inc., 2019) and R (Version 4.0.0; R Core Team, 2017) functions (https://osf.io/ypmdj). For the RSA analyses irrespective of memory, we modelled dissimilarities between all item pairs, treating all trials in the same way (see Fig. 3C).

##### fMRI data RDMs

Parameter estimates were extracted from gray matter voxels in each ROI for all trials of interest. For each voxel, these betas were then normalized by dividing them by the standard deviation of its residuals (Walther et al., 2016). As for the model RDMs, we constructed separate fMRI data RDMs for the true and false subsequent memory and overall object processing analyses. For the true subsequent memory analysis, the fMRI data RDM represented activity patterns for concepts subsequently tested as old, and for the false subsequent memory analysis, the fMRI data RDM represented activity patterns for concepts subsequently tested as lures. For the overall analysis, the RDM represented activity patterns for all study trials. For the fMRI data RDMs for the subsequent memory analysis, as for the model RDMs, we computed dissimilarity between all pairings of subsequently remembered (or falsely recognized) items, and between all pairings of subsequently forgotten (or correctly rejected) items, omitting pairings between different trial types. Distance between each item pair was computed as 1 - Pearson’s correlation, creating a dissimilarity matrix.

##### Fitting model to data RDMs

Each fMRI data RDM was compared with each theoretical model RDM using Spearman’s rank correlation, and the resulting dissimilarity values were Fisher-transformed. For the subsequent memory analysis, we tested for significant positive and negative similarities between model RDM and fMRI data RDMs at the group level using a two-sided Fisher’s one-sample randomization (10,000 permutation) test for location with a Bonferroni correction over 6 ROIs. The permutation distribution of the test statistic *T* enumerates all the possible ways of permuting the correlation signs, positive or negative, of the observed values and computes the resulting sum. Thus, for a two-sided hypothesis, the *p*-value is computed from the permutation distribution of the absolute value of *T*, calculating the proportion of values in this permutation distribution that are greater or equal to the observed value of *T* (Millard and Neerchal, 2001). For the overall analysis we only tested for significant positive similarities between model RDM and fMRI data RDMs (Clarke and Tyler, 2014), using a one-sided test, in which the *p*-value is evaluated as the proportion of sums in the permutation distribution that are greater than or equal to the observed sum *T* (Millard and Neerchal, 2001). To find the unique effect of model RDMs, each fMRI data RDM showing a significant effect was also compared with each theoretical model RDM while controlling for effects of all other significant model RDMs (using partial Spearman’s rank correlations).

#### RSA searchlight analysis

In addition to the targeted ROI analysis, we ran a whole-brain searchlight analysis. This followed the same 3 main steps as the ROI analysis (see RSA region of interest analysis). For each voxel, the fMRI data RDM was computed from parameter estimates for gray matter voxels within a spherical searchlight of radius 7 mm, corresponding to maximum dimensions 5 × 5 × 5 voxels. Dissimilarity was again estimated using 1 - Pearson’s correlation. As in the ROI analysis, this fMRI data RDM was compared with the model RDMs, and the resulting dissimilarity values were Fisher transformed and mapped back to the voxel at the center of the searchlight. The similarity map for each model RDM and participant was then normalized to the MNI template space (see Image preprocessing). For each model RDM, the similarity maps were entered into a group-level random-effects analysis and thresholded using permutation-based statistical nonparametric mapping (SnPM; http://www.nisox.org/Software/SnPM13/). This corrected for multiple comparisons across voxels and the number of theoretical model RDMs. As for the ROIs we performed two-tailed tests in the subsequent memory analyses and one-tailed tests for the overall analysis. Variance smoothing of 6 mm FWHM and 10,000 permutations were used in all analyses. We used cluster-level inferences with FWE-correction at *α* = 0.025 in each direction for the two-tailed tests and *α* = 0.05 for the one-tailed test, in both cases with a cluster forming threshold of 0.005 uncorrected. All results are presented on an inflated representation of the cortex using the BrainNet Viewer (Xia et al., 2013, http://www.nitrc.org/projects/bnv/) based on a standard ICBM152 template.

### Univariate fMRI analysis

In addition to RSA, we used univariate analysis to test whether activation in PrC was related to the conceptual confusability of an object, in a replication of Clarke and Tyler (2014), and whether this activation predicted memory. We also used activations to define additional ROIs (see Regions of interest). The first level GLM for each participant included one regressor of interest for each of the 4 experimental conditions (subsequent hits, misses, false alarms, and correct rejections). For each condition, we also included 4 linear parametric modulator regressors representing concept confusability values for each concept with other concepts in the CSLB property norms (Devereux et al., 2014). We first computed a semantic similarity score between each pair of concepts (see RSA region of interest analysis, Model RDMs). The concept confusability score of each concept was then equal to the sum of squared similarities between it and the other concepts in the set. This was equivalent to a weighted sum of pair-wise similarities in which each weight was the between-concept similarity itself, a measure used in our recent behavioral study (Naspi et al., 2020). As also specified in the preregistration, since the results of the concept confusability analysis diverged from those of Clarke and Tyler (2014), we ran an additional analysis using a measure of concept confusability with a stronger weighting scheme equivalent to theirs. They defined concept confusability as the exponential of the ranked similarities of all the paired concepts, which is very close to a nearest neighbor scheme in which each concept’s similarity is equal to its similarity to the most similar concept in the set. Due to our larger number of items the exponential weighting produced extremely large weights, so we substituted the simpler nearest neighbor scheme (the two measures were correlated at r = 0.98). We used an explicit mask including only voxels which had at least a 0.2 probability of being in grey matter as defined using the MNI template. To permit inferences about encoding condition effects across participants, contrast images were submitted to a second-level group analysis (one sample *t-*test) to obtain *t*-statistic maps. The maps were thresholded at *p* < 0.05, FWE-corrected for multiple comparisons at the voxel level using SPM (the preregistration specified 3dClustSim in AFNI, but this function had since been updated (Cox et al., 2017) so for simplicity we used the SPM default). Only regions whose activations involved contiguous clusters of at least 5 voxels were retained as ROIs for subsequent RSA analysis.

### Code accessibility

All analyses were performed using custom code and implemented either in MATLAB or R. All code and the data for the behavioral and the fMRI analyses are available through https://osf.io/z4c62/.

## Results

### Memory task performance

In the study phase, participants correctly identified most of the time whether concepts began with a consonant or vowel on the incidental encoding task (M proportion = 0.78). Analysis on task engagement (see Materials and Methods, Behavioral data) using a GLMM showed that accuracy at encoding did not differ according to whether items that were tested as studied were later remembered relative to forgotten (*β* = 0.110, SEM = 0.242, *z* = 0.456, *p* = 0.649), or whether items that were tested as lures were later falsely recognized relative to correctly rejected (*β* = 0.051, SEM = 0.202, *z* = 0.251, *p* = 0.802). Similarly, a linear mixed model did not reveal any difference in RTs related to subsequent old items that were later remembered relative to forgotten (*β* = 0.002, SEM = 0.017, *t* = 0.123, *p* = 0.902), or subsequent lures that were later falsely recognized relative to correctly rejected (β = -0.013, SEM = 0.015, *t* = - 0.873, *p* = 0.383). Thus, the fMRI subsequent memory effects are not attributable to differences in accuracy or time on task at encoding.

At test, as a simple check on the overall level of performance we used the discrimination index *Pr*, i.e., the difference between the probability of a hit to studied items and the probability of a false alarm to novel items. All participants passed the preregistered inclusion criterion of *Pr* > 0.1. Overall, discrimination collapsed across confidence was very good (M = 0.649, SD = 0.131, *t*_(27)_ = 26.259, *p* < 0.001). Discrimination was also above chance for high confidence (M = 0.771, SD = 0.152, *t*_(27)_ = 26.868, *p* < 0.001) and low confidence judgments (M = 0.330, SD = 0.145, *t*_(27)_ = 12.014, *p* < 0.001). This suggests that low confidence responses at test carried veridical memory, so we followed our preregistered plan to include trials attracting both high and low confidence responses in the subsequent memory analysis. Following an analogous procedure for false recognition of similar lures corrected by subtracting the proportion of false alarms to novel items, we also found that this was significantly above chance for judgments collapsed across confidence (M = 0.271, SD = 0.090, *t*_(27)_ = 15.996, *p* < 0.001), and for both high confidence (M = 0.293, SD = 0.133, *t*_(27)_ = 11.618, *p* < 0.001) and low confidence (M = 0.157, SD = 0.160, *t*_(27)_ = 5.187, *p* < 0.001) considered separately.

We then used a GLMM to quantify the influence of perceptual and semantic variables on memory performance according to item status. Our variables of interest were condition (studied, lure, or novel), concept confusability, C1 visual confusability, and color confusability (see Behavioral data for details). Results revealed modulations of memory by perceptual and semantic variables in line with our recent behavioral study (Naspi et al., 2020). People were less likely to recognize studied items for which the low-level visual representations (C1) were more similar to those of their nearest neighbor (*β* = -0.166, SEM = 0.064, *z* = -2.584, *p* = 0.015), and also less likely to recognize studied items with high concept confusability relative to novel items (*β* = -0.533, SEM = 0.067, *z* = -7.963, *p* < 0.001). As expected, concept confusability also had a substantial effect on false recognition of similar lures relative to novel items, whereby images whose concepts were more confusable with other concepts in the set were less likely to be falsely recognized (*β* = -0.273, SEM = 0.064, *z* = -4.292, *p* < 0.001).

### Preregistered RSA analysis in regions of interest

#### Perceptual and semantic representations predict true recognition

To examine representations engaged during successful encoding we compared the fit of early visual, color, animal-nonbiological-plant, and semantic feature models for studied items tested as old that were subsequently remembered (number of trials, M = 61.41; range = 60-146) versus forgotten (number of trials, M = 19.93; range = 17-104) (Fig. 4A). These comparisons were bidirectional, since engagement of perceptual and/or semantic processing in a region might either support or be detrimental to later memory. Thus, we used a two-sided Fisher’s randomization test *T*. In posterior ROIs, engagement of both perceptual and finer-grained semantic representations tended to predict successful later recognition. In EVC, the early visual model strongly predicted later true recognition of studied items (M = 0.07, 95% CI [0.05, 0.09], *T* = 1.86, *p* < 0.001). Thus, when the neural patterns at study were representing visual information, items were more likely to be correctly recognized. Both the early visual and semantic feature models also predicted true recognition in pVTC (M = 0.03, 95% CI [0.02, 0.04], *T* = 0.82, *p* < 0.001, and M = 0.02, 95% [CI: 0.01, 0.04], *T* = 0.67, *p* = 0.007, respectively). In contrast, taxonomic semantic representations coded more anteriorly were associated with later forgetting. In aVTC and in the LIFG, model fit for categorical semantic information represented by the animal-nonbiological-plant domain was less for remembered than forgotten studied items (M = -0.01, 95% CI [-0.02, -0.01], *T* = 0.35, *p* = 0.001, and M = -0.02, 95% CI [-0.04. -0.01], *T* = 0.65, *p* = 0.004, respectively). Thus, when neural patterns in these regions were aligned with items’ taxonomic categories, participants were less likely to successfully recognize them. No other results were significant.

**Figure 4.**
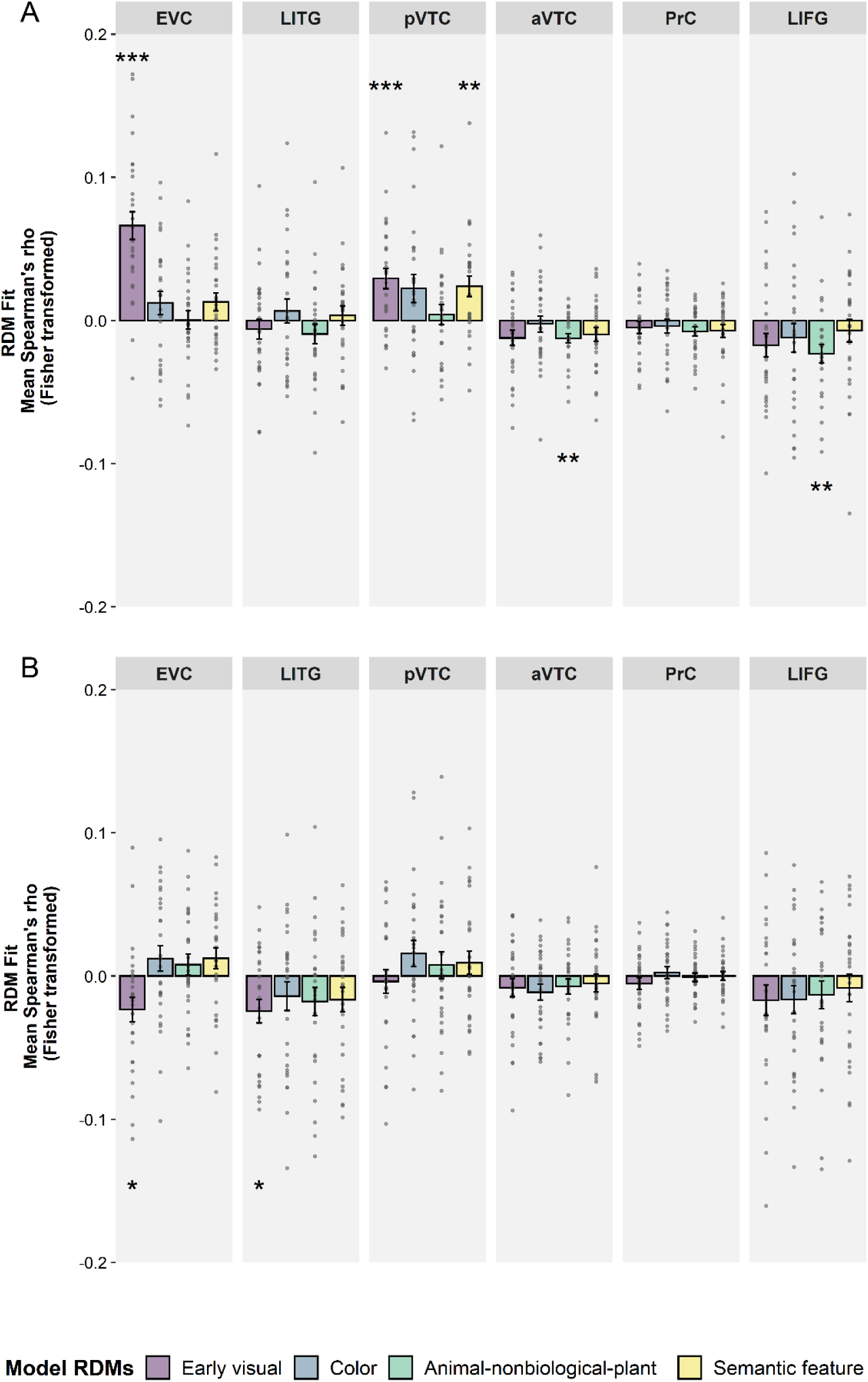
Perceptual and semantic representations predicting subsequent memory for *a priori* ROIs and models. Plots show the group mean Fisher-transformed Spearman correlation coefficient reflecting perceptual and semantic representations associated with: **A**, true subsequent memory; **B**, false subsequent memory. Error bars represent the standard error of the mean (SEM) across participants. Asterisks indicate models for which Spearman’s rho differed significantly from zero at the group level (two-sided Fisher’s randomization test for location; Bonferroni correction calculated multiplying the uncorrected *p*-value by the number of comparisons made). * *p* < 0.05, ** *p* < 0.01, *** *p* < 0.001

We also checked which representations showed unique effects that predicted memory after controlling for effects of other significant models using partial correlation. In pVTC, only the early visual model uniquely predicted successful recognition memory for studied items (M = 0.02, 95% CI [0.01, 0.03], *T* = 0.64, *p* = 0.004) (but see Exploratory ROI analysis).

#### Weak perceptual representations predict false recognition

To examine how the perceptual and semantic representations embodied in our theoretical models contributed to subsequent memory for lures, we compared RSA model fit for items that were later falsely recognized (number of trials, M = 30.71; range = 26-107) versus correctly rejected (number of trials, M = 50.61; range = 54-131) (Figure 4B). In posterior regions, weaker low-level visual representations of pictures predicted subsequent false recognition of lures. We observed this pattern in both the EVC and the LITG (M = -0.02, 95% CI [-0.04, -0.01], *T* = 0.66, *p* = 0.047, and M = -0.02, 95% CI [-0.04, -0.01], *T* = 0.69, *p* = 0.026, respectively). Thus, when neural patterns in these regions were not aligned with the early visual model, items were more likely to be falsely recognized. No other results were significant.

#### Perceptual and semantic object processing irrespective of memory

Replicating Clarke and Tyler (2014), we also examined the perceptual and semantic representations of objects that were reflected in fMRI activity patterns regardless of memory encoding. The results (Fig. 5) showed that while visual information is broadly represented posteriorly, activity patterns in the aVTC, PrC, and LIFG reflect finer-grained semantic information. Posteriorly, EVC showed a strong relationship with the low-level visual model (M = 0.08, 95% CI [0.06, 0.10], *T* = 2.21, *p* < 0.001), and a weaker but significant relation with the semantic feature model (M = 0.01, 95% CI [0.00, 0.01], *T* = 0.20, *p* = 0.032). More anteriorly, the low-level visual and semantic feature models were both significantly related to activity patterns in pVTC (M = 0.04, 95% CI [0.03, 0.04], *T* = 1.00, *p* < 0.001, and M = 0.02, 95% CI [0.02, 0.03], *T* = 0.60, *p* < 0.001, respectively) and in LITG (M = 0.01, 95% CI [0.00, 0.02], *T* = 0.26, *p* < 0.038, and M = 0.02, 95% CI [0.01, 0.02], *T* = 0.45, *p* < 0.001, respectively). At the apex of the ventral visual pathway, semantic feature information was coded in both the bilateral aVTC (M = 0.01, 95% CI [0.00, 0.01], *T* = 0.17, *p* = 0.006) and in bilateral PrC (M = 0.01, 95% CI [0.00, 0.01], *T* = 0.19, *p* < 0.001). These findings replicated those of Clarke and Tyler (2014). The specific semantic properties of objects were also represented in the LIFG (M = 0.01, 95% CI [0.01, 0.02], *T* = 0.30, *p* = 0.001).

**Figure 5.**
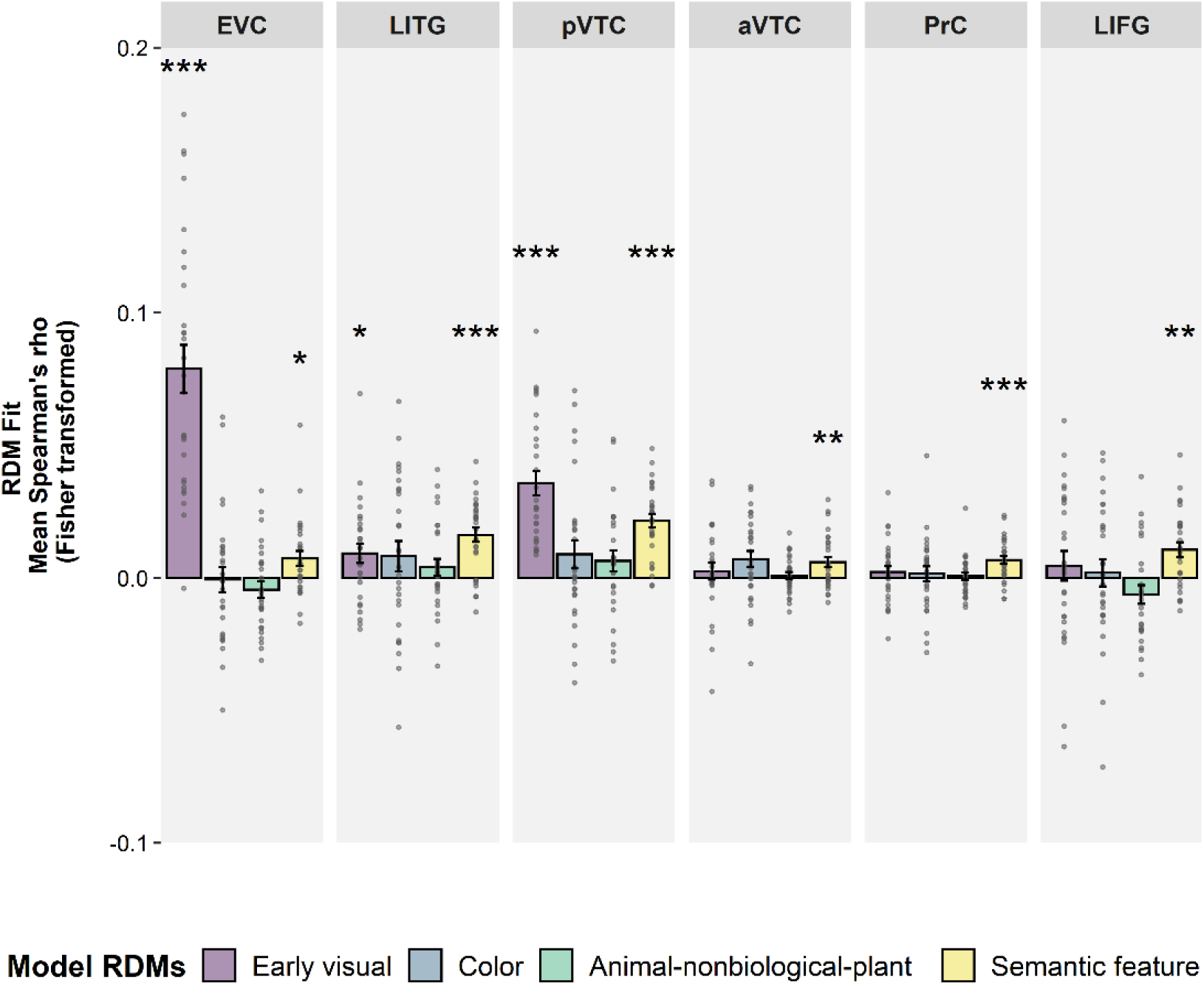
Semantic and perceptual representations represented in ROIs regardless of memory encoding. Region of Interest (ROI) results comparing four model RDMs to patterns of activity along the ventral stream and frontal regions. Error-bars are standard error of the mean (SEM) across subjects. Asterisks above and below the bars depict *p*-values for tests of whether each individual Spearman’s correlation is greater than zero (one-sided Fisher’s randomization test for location; Bonferroni correction calculated multiplying the uncorrected *p*-value by the number of comparisons made). * *p* < 0.05, ** *p* < 0.01, *** *p* < 0.001

We then ran a partial correlation on those ROIs showing significant effects for different models. As expected, patterns of activity in the EVC were uniquely related to the early visual model (M = 0.08, 95% CI [0.06, 0.10], *T* = 2.20, *p* < 0.001), replicating Clarke and Tyler’s (2014) results. Thus, the semantic feature model was no longer significant when the early visual model was controlled for. More anteriorly, both low-level visual and semantic feature information were uniquely related to the pattern of activity in the pVTC (M = 0.03, 95% CI [0.03, 0.04], *T* = 0.96, *p* < 0.001, and M = 0.02, 95% CI [0.01, 0.02], *T* = 0.54, *p* < 0.001, respectively). However, after controlling for the low-level visual model, activity patterns in the LITG were only uniquely associated with semantic feature representations (M = 0.02, 95% CI [0.01, 0.02], *T* = 0.44, *p* < 0.001). Thus, like Clarke and Tyler (2014), we found that visual information is represented in early visual regions. We also replicated their finding that semantic feature similarity information was coded more anteriorly in the PrC, and found further, also anterior, regions that showed a similar pattern, in the aVTC and the LIFG (see also RSA searchlight fMRI analysis).

### Exploratory RSA analysis in regions of interest

Perceptual and semantic representations in pVTC subdivisions predict true recognition In the preregistered analyses reported above, our large pVTC ROI showed evidence of both visual and semantic feature representations predicting memory success. We therefore explored whether four subdivisions of this large bilateral region showed distinct effects: the LG, ITG, FG, and PHC (see Regions of interest). Moreover, given our strong *a priori* prediction of involvement of PrC in subsequent memory, we ran exploratory analyses in left and right PrC separately. The results are shown below in Figure 6. Posteriorly, in bilateral LG, perceptual information related to the early visual model predicted later recognition of studied items (M = 0.03, 95% CI [0.01, 0.04], *T* = 0.74, *p* = 0.002), as it did in the EVC ROI. In contrast, more anteriorly, activity patterns in the FG related to both the low-level visual and semantic feature models predicted subsequent true recognition (M = 0.03, 95% CI [0.02, 0.05], *T* = 0.87, *p* = 0.002, and M = 0.04, 95% CI [0.02, 0.05], *T* = 1.01, *p* < 0.001, respectively), as did categorical semantic information represented by the animal-nonbiological-plants model in the PHC (M = 0.02, 95% CI [0.01, 0.03], *T* = 0.55, *p* = 0.019). Lastly, activity related to the categorical semantic model in the left PrC predicted subsequent forgetting (M = -0.01, 95% CI [-0.02, 0.00], *T* = 0.28, *p* = 0.023).

**Figure 6.**
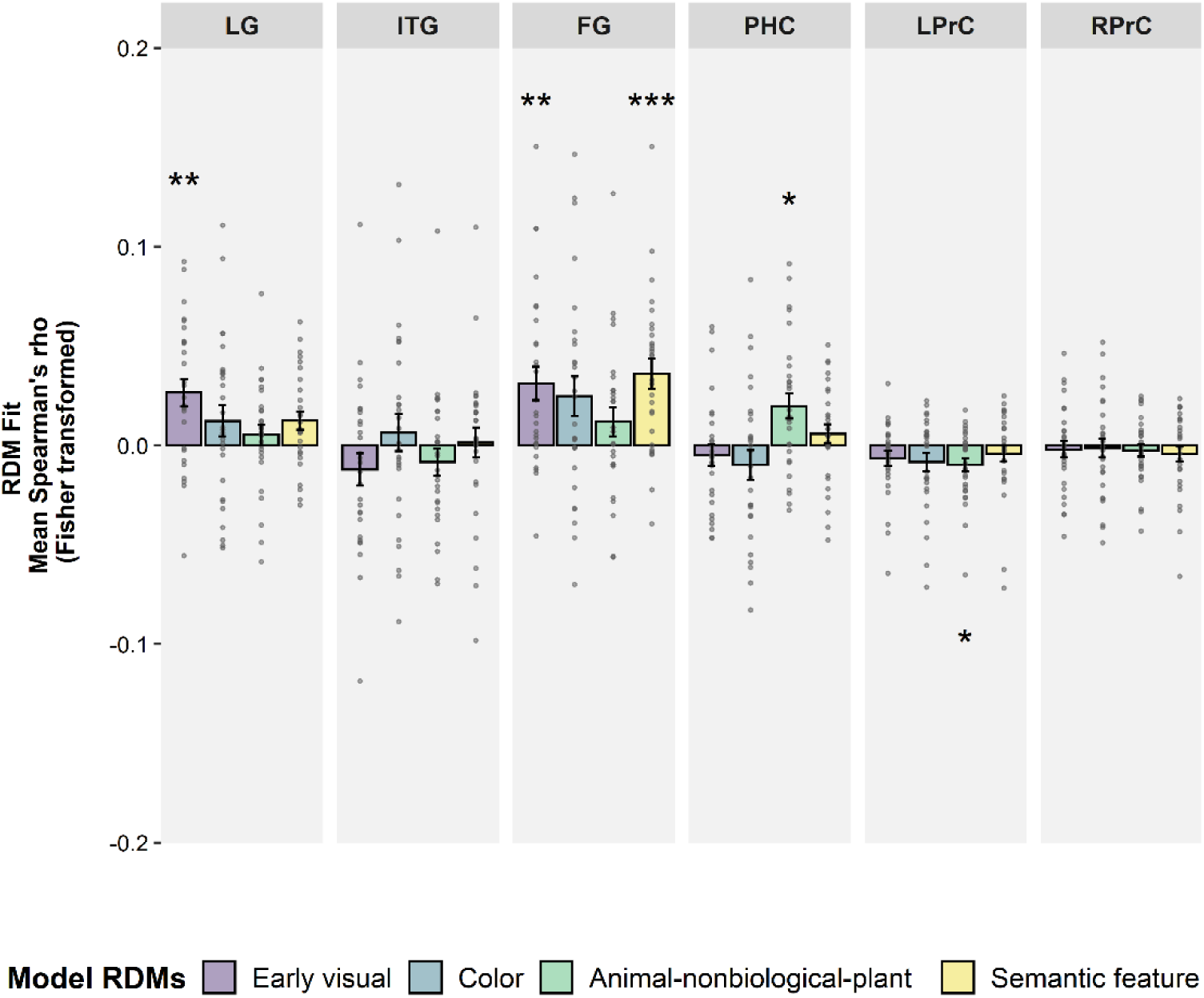
Perceptual and semantic representations predicting true subsequent memory in exploratory ROIs. Plots show the group mean Fisher-transformed Spearman correlation coefficient reflecting perceptual and semantic representations associated with true subsequent memory. Error bars represent the standard error of the mean (SEM) across participants. Asterisks indicate significance of tests of group level differences of Spearman’s rho from zero (two-sided Fisher’s randomization test for location; Bonferroni correction calculated multiplying the uncorrected *p*-value by the number of comparisons made). * *p* < 0.05, ** *p* < 0.01, *** *p* < 0.001

A partial correlation analysis for the FG (which showed effects of multiple models) confirmed that both the early visual and semantic feature models were uniquely associated with later true recognition (M = 0.02, 95% CI [0.01, 0.03], *T* = 0.58, *p* = 0.034, and M = 0.03, 95% CI [0.01, 0.04], *T* = 0.71, *p* = 0.002, respectively). Thus, both simple visual and object-specific semantic information contributed to memory after controlling for each other.

Lastly, following our main analyses of true and false memory encoding, we wanted to check for evidence that some of the key results differed according to encoding type. Thus, we compared the fit of our theoretical models for studied items tested as old that were subsequently remembered versus lures that were subsequently falsely recognized. Results showed that low-level visual information mapped in EVC was stronger for items that were subsequently remembered than falsely recognized (M = 0.04, 95% CI [0.02, 0.06], *T* = 1.16, *p* < 0.001). No other results were significant at the Bonferroni-corrected threshold, but without a correction the theoretically important object-specific semantic representations in FG were also stronger for true than false recognition (M = 0.02, 95% CI [0.00, 0.04], *T* = 0.61, *p* = 0.030).

### Preregistered RSA searchlight analysis

#### Perceptual and semantic representations associated with memory encoding

The RSA searchlight analysis tested for any further brain regions coding for perceptual and semantic information associated with memory encoding (Figure 7 and Table 1). The true subsequent memory models showed significant fit to activity patterns in several areas beyond the *a priori* ROIs. The color similarity model was related to patterns in the right parietal opercular cortex, superior frontal gyrus, and precentral gyrus, and this representation at encoding predicted later successful recognition of studied items. Fine-grained semantic features represented in the right lateral occipital cortex (LOC) also predicted true recognition. Coarse categorical semantic representations in right inferior frontal gyrus (RIFG; BA44/45/47) and frontal pole (FP) were associated with later forgetting, paralleling the findings for the *a priori* ROI in LIFG (BA44/45).

**Figure 7.**
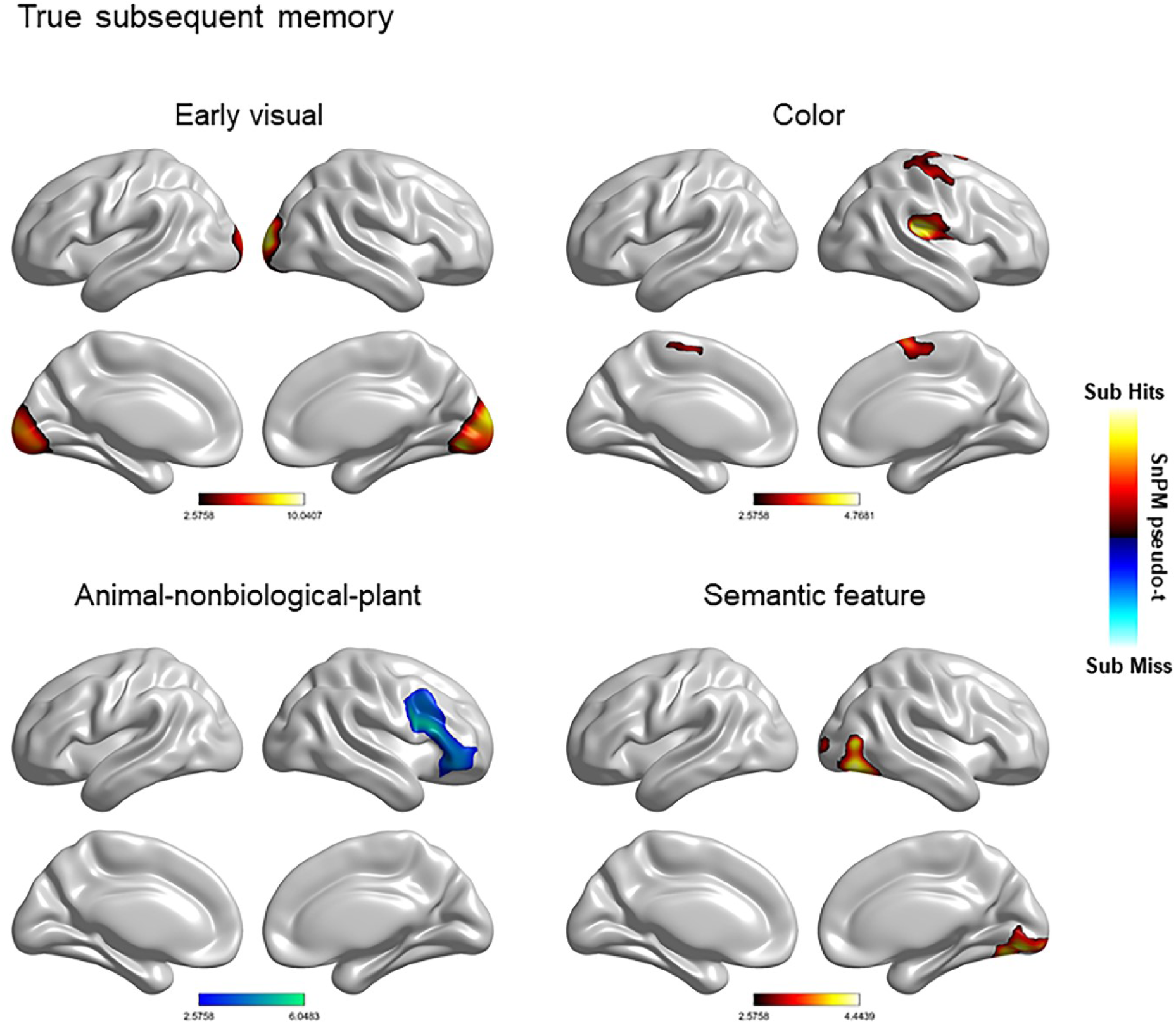
RSA searchlight results for perceptual and semantic models. The figure shows regions in which multivoxel activity pattern predicted successful subsequent true recognition (hot map) and unsuccessful true recognition (i.e., subsequent forgetting, cool map). All significant clusters are shown at the FWE-corrected threshold used for analysis (see Materials and Methods: RSA searchlight analysis). No suprathreshold voxels survived for the subsequent false recognition models. Similarity maps are presented on an inflated representation of the cortex based on the normalized structural image averaged over participants.

**Figure 8.**
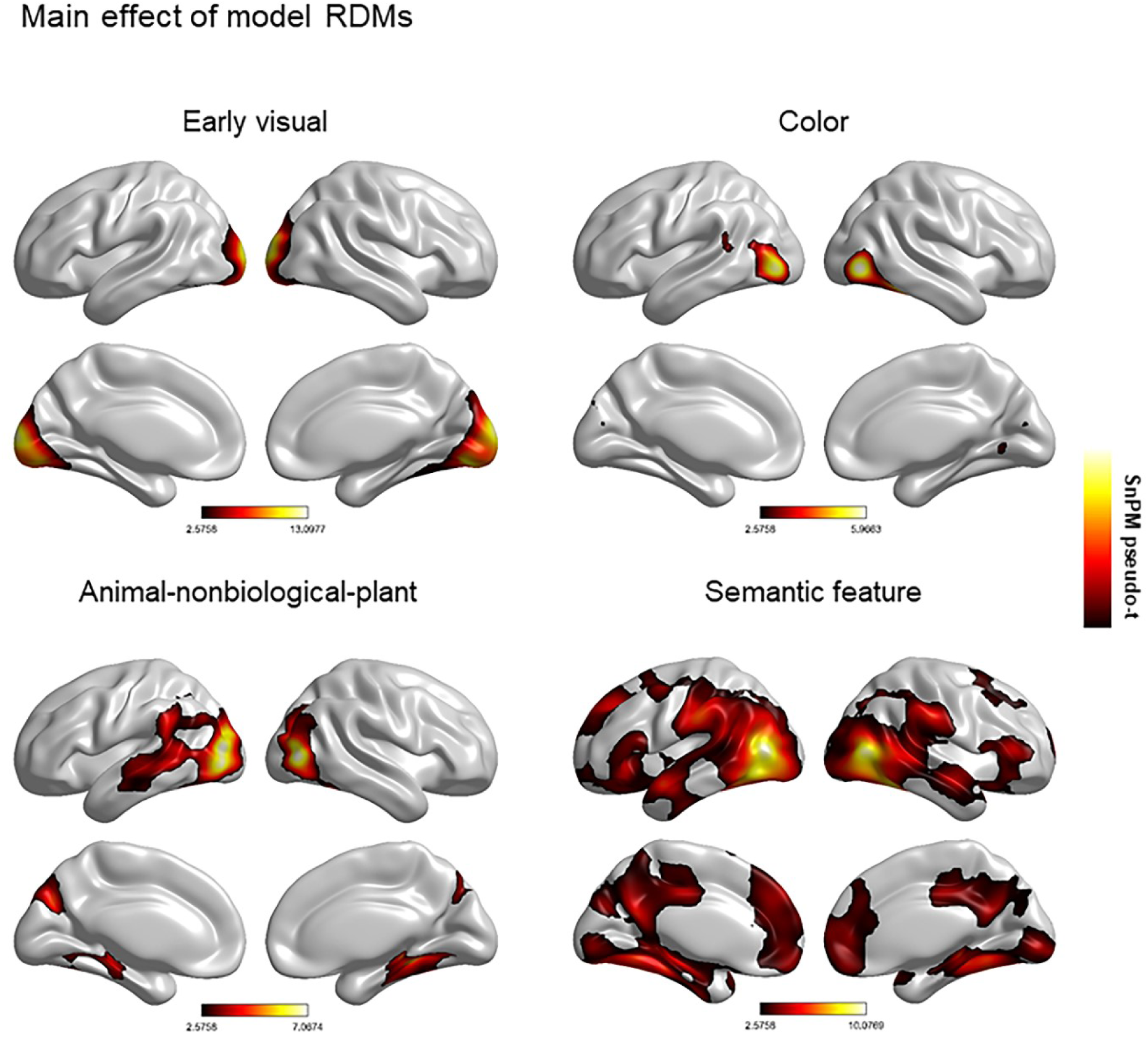
RSA searchlight results for perceptual and semantic models. The figure shows regions in which multivoxel activity pattern was associated with object processing (i.e., irrespective of memory encoding). All significant clusters are shown at the FWE-corrected threshold used for analysis (see Materials and Methods: RSA searchlight analysis). Similarity maps are presented on an inflated representation of the cortex based on the normalized structural image averaged over participants.

**Table 1.**
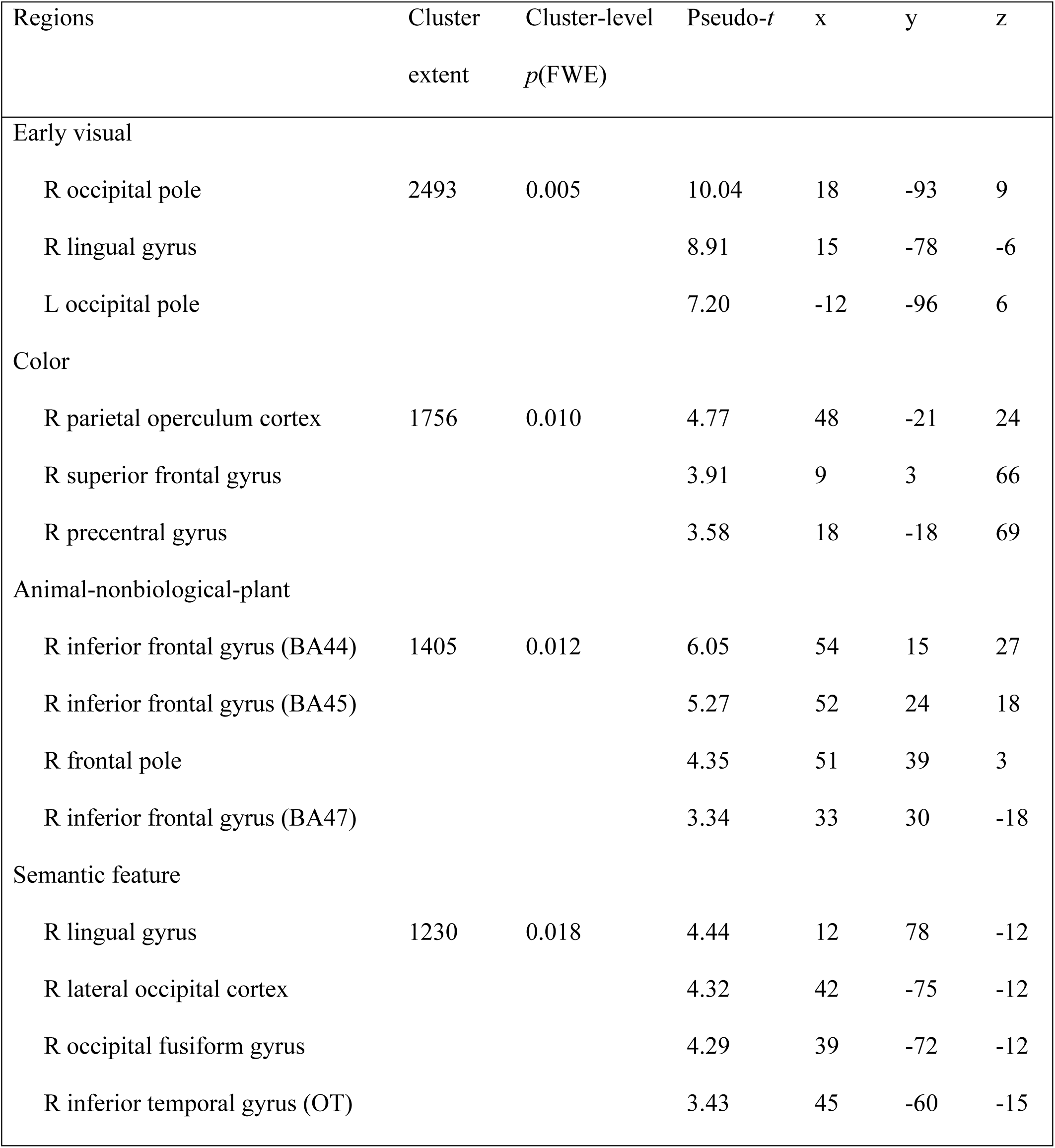
RSA searchlight results showing perceptual and semantic effects on successful true memory encoding

MNI coordinates and significance levels are shown for the peak voxel in each cluster. Anatomical labels are provided for peak locations in each cluster. Effects in clusters smaller than 20 voxels not shown. OT = Occipito-temporal division.

#### Perceptual and semantic object processing irrespective of memory

Searchlight analysis was also conducted for the perceptual and semantic model RDMs across all trials regardless of memory encoding (Fig. 5 and Table 2). The models showed significant fit to multivoxel activity patterns in several areas beyond the *a priori* ROIs. In particular, the effects for the color model were largely restricted to the right lateral occipital cortex, right middle temporal gyrus, and intracalcarine cortex, but also extended into the left lateral occipital cortex and supramarginal gyrus. Categorical semantic representations represented by the animal-nonbiological-plant domain were largely restricted to posterior parts of the ventral stream, highlighting the coarse nature of object information represented in the posterior ventral temporal cortex. This included the right temporal fusiform cortex, the right lingual gyrus, and the posterior division of parahippocampal cortex, but also extended into the middle temporal lobe. In contrast, representation of finer-grained semantic properties of objects extended more anteriorly in the ventral pathway beyond the preregistered ROIs, into bilateral hippocampus, temporal pole and ventromedial frontal regions.

**Table 2.**
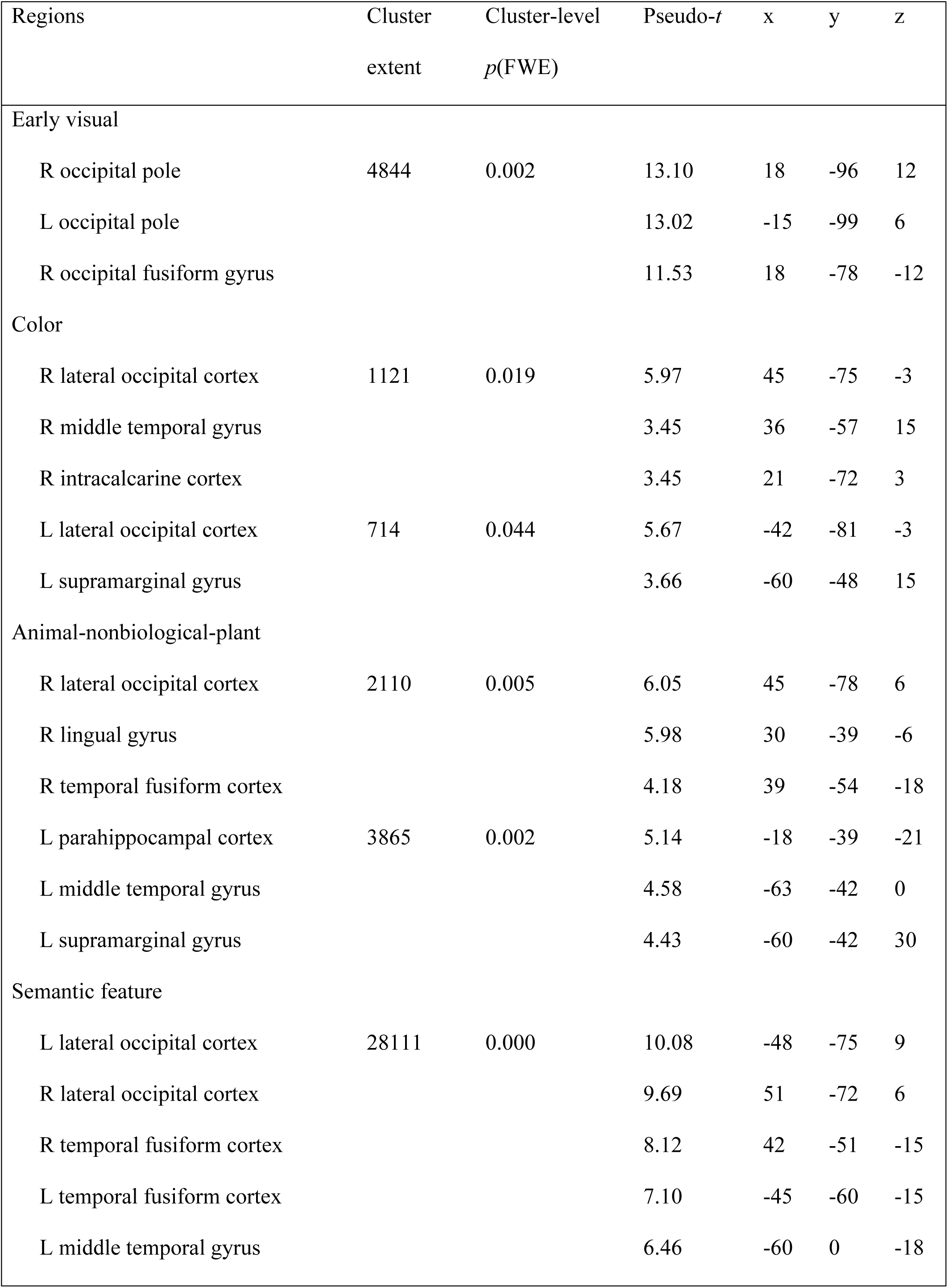
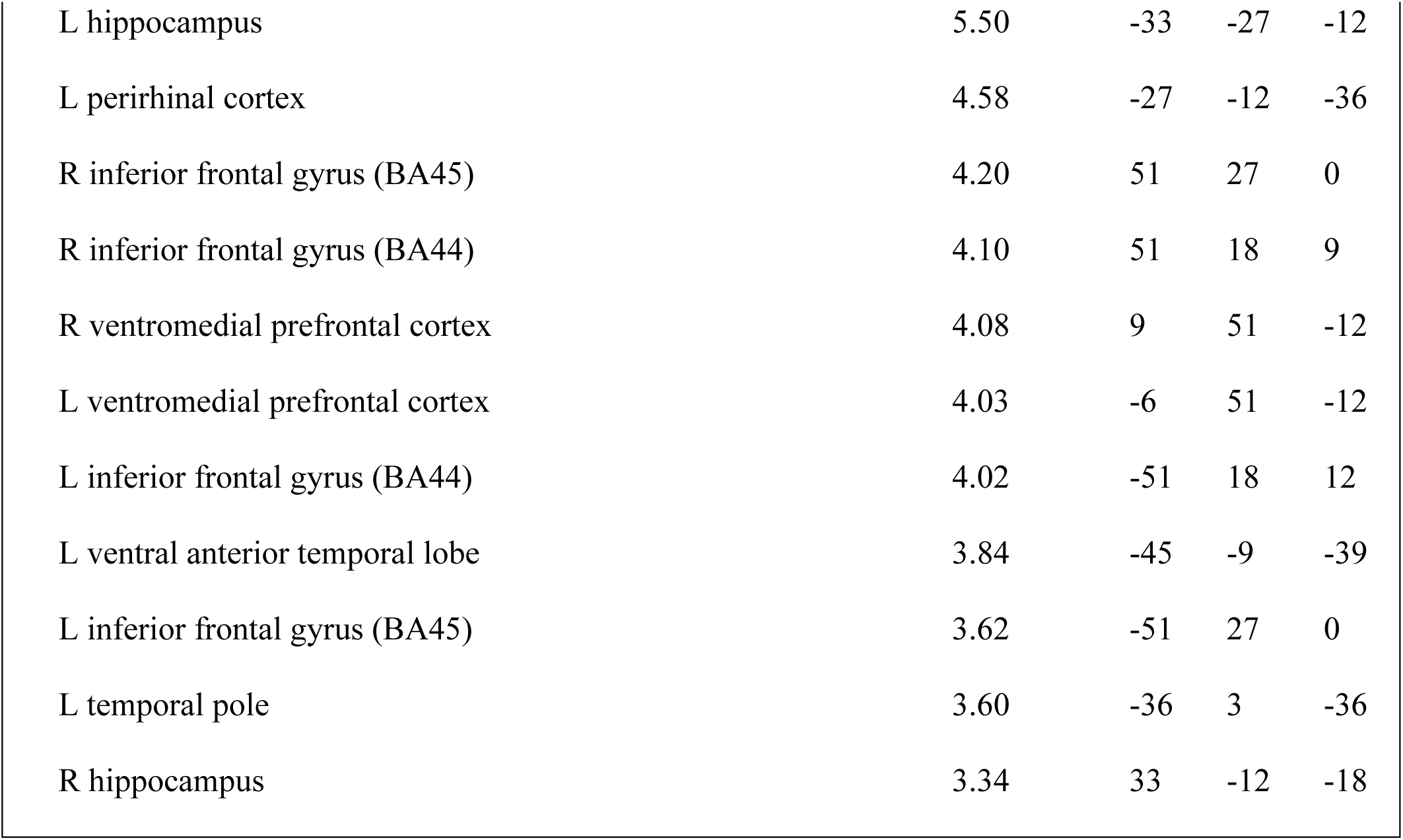
RSA results showing perceptual and semantic effect of object processing

MNI coordinates and significance levels shown for the peak voxel in each cluster. Anatomical labels are provided for locations in each cluster. Effects in clusters smaller than 20 voxels not shown.

### Preregistered univariate fMRI analysis

#### Encoding activity predicting true and false recognition

Univariate analysis was run to derive ROIs for RSA based on subsequent memory effects in regions where prior literature is suggestive, but not clear, regarding their involvement. This showed significant activation for subsequently remembered > subsequently forgotten items in the LITG (cluster size: *k* = 13, *p* < 0.05 FWE). No significant activation was revealed for subsequently falsely recognized > subsequently corrected rejected items after FWE correction.

#### Parametric effect of concept confusability

Finally, we were interested in the specific role of the PrC, and possibly aVTC, in processing conceptually confusable objects. These regions were not related to parametric changes in concept confusability regardless of memory encoding. Therefore, we did not replicate Clarke and Tyler (2014)’s finding of increased activation for more conceptually confusable objects (uncorrected *p* = 0.139 and *p* = 0.05 for PrC and aVTC, respectively). Subsequent memory effects were also not significant at the preregistered FWE-corrected threshold. However, at an uncorrected threshold, activity associated with concept confusability was greater for subsequently forgotten than remembered items in right PrC (cluster size: *k* = 12, *p* < 0.005) and bilateral aVTC (right cluster size: *k* = 19, *p* < 0.001; left cluster size: *k* = 6, *p* < 0.001). Activity associated with concept confusability was also greater for subsequently falsely recognized than correctly rejected items in bilateral PrC (right cluster size: *k* = 35, *p* < 0.005; left cluster size: *k* = 11, *p* < 0.005), and right aVTC (cluster size: *k =* 22, *p* < 0.005), and for subsequently falsely recognized than remembered items in bilateral PrC (right cluster size: *k* = 25, *p* < 0.005; left cluster size: *k* = 12, *p* < 0.005), and righy aVTC (cluster size: *k* = 16, *p* < 0.005).

## Discussion

Our results show that semantic and perceptual representations play distinct roles in true and false memory encoding. By combining explicit models of prior conceptual knowledge and image properties with a subsequent memory paradigm, we were able to probe their separate contributions to encoding of objects. Fine-grained perceptual and semantic processing in the ventral visual pathway both predicted later recognition of studied objects, while coarser-grained categorical semantic information processed more anteriorly predicted forgetting. In contrast, only weak low-level visual representations in posterior regions predicted false recognition of similar objects. The data provide the first direct tests of fuzzy-trace theory’s assumptions about how memories are encoded, and suggest that semantic representations may contribute to specific as well as gist memory phenomena (Brainerd and Reyna, 2002).

Our results for the early visual model converge with studies showing univariate subsequent memory effects in the same regions (Kim and Cabeza, 2007; Kirchhoff et al., 2000; Pidgeon and Morcom, 2016; Wagner et al., 1998). Distributed low-level visual representations in early visual cortices predicted successful later recognition of specific studied objects. The C1 HMax representations embody known properties of primary visual cortex relating to local edge-orientations in images (Kamitani and Tong, 2005), and this model clustered our object images by overall shape and orientation (Fig. 2). These results converge with Davis et al. (2020)’s recent finding that RSA model fit for an early layer of a deep convolutional neural network (DNN) in early visual cortex predicted later memory for pictures. Our models are directly interpretable, allowing us to show unambiguously that representing lower-level properties available in the presented images contributes to memory.

In late visual regions, such as LG and FG, activity patterns fitting the early visual model also predicted true recognition (Fig. 5 and 7), as hypothesized based on activation studies (Garoff et al., 2005; Kim, 2011; Kirchhoff et al., 2000; Stern et al., 1996; Vaidya et al., 2002). We also found that specific object features coded in FG predicted true recognition. These pVTC regions receive low-level properties as input to compute complex shape information (Kanwisher, 2001). Emerging data suggest that the FG supports visuo-semantic processing of modality-specific semantic features. Devereux et al. (2018) combined deep visual and semantic attractor networks to model the transformation of vision to semantics, revealing a confluence of late visual representations and early semantic feature representations in FG (see also Tyler et al., 2013). This converges with Martin et al.’s (2018) finding that FG activity reflected representations of explicitly rated visual object features. Davis et al. (2020) reported that in FG the mid-layer of a visual DNN predicted memory for object names when the objects were forgotten, while semantic features of the object images predicted memory for the images when the names were forgotten. Our findings clarify that both image-based visual codes and non-image-based semantic feature codes are represented during successful encoding. Together, the data further suggest that this initial extraction of semantic features from vision is important for the effective encoding of memories of specific objects, more than false recognition of similar objects.

More anteriorly, taxonomic categorical representations in aVTC and left PrC predicted forgetting of studied items. Similar findings in LIFG support the idea that coarse-grained domain-level semantic processing is detrimental to memory for specific objects. LIFG typically shows strong univariate subsequent memory effects for verbal or nameable object stimuli (Kim, 2011). It is thought to support selection and control processes involved in elaborative semantic encoding (Jackson et al., 2015; Prince et al., 2007). Our overall analysis showed that object-specific semantic information was represented in this region, but did not predict recognition. One possibility is that domain-level taxonomic processing impeded selection of specific semantic information. Another possibility, in line with the levels of processing principle, is that the object naming encoding task did not strongly engage semantic control operations that promote subsequent memory (Craik and Lockhart, 1972; Otten and Rugg, 2001). Object naming depends on basic-level object-specific processing in the FG, consistent with the current findings (Taylor et al., 2012). Future studies can test this by manipulating cognitive operations at encoding to determine whether the representations promoting later memory are also task-dependent.

The absence of any association between object-specific representations in PrC and encoding was unexpected, although we replicated Clarke and Tyler (2014)’s central finding that PrC represents object-specific semantic features. The PrC encodes complex conjunctions of visual (Barense et al., 2012; Bussey et al., 2002) and semantic features (Bruffaerts et al., 2013; Clarke and Tyler, 2014) that enable fine-grained object discrimination and may contribute to later item memory (Brown and Aggleton, 2001; Yonelinas et al., 2005). As the object-specific semantic model fit embodied both shared and distinctive feature information, we ran a further, univariate analysis to examine the directional effect of shared features (concept confusability). We did not replicate Clarke & Tyler’s (2014) finding that PrC activation was higher overall for more confusable objects, interpreted in terms of feature disambiguation. However, we found preliminary evidence that in both PrC and vATL, activity correlating with concept confusability predicted forgetting of studied objects. This is consistent with our finding that concept confusability strongly impairs true recognition, as well as discrimination between studied objects and lures (Naspi et al., 2020), results replicated here. The RSA data also suggest an interpretation of Davis et al.’s (2020) report that semantic feature model fit in PrC predicted later true recognition of object concepts when their pictures were forgotten, which may correspond to nonspecific encoding.

An important and novel feature of our study is the investigation of the representational content associated with encoding of false memories. Our results revealed that weak visual representations coded in EVC and extending to LITG predicted later false recognition (Fig. 5), and model fit differed significantly from true recognition. This supports fuzzy-trace theory’s proposal that visual detail is encoded in specific memory traces that confer robustness to later true recognition (Brainerd and Reyna, 2002). Several univariate fMRI studies of memory retrieval have shown greater early and late visual cortex activation for true than false memories of objects (Dennis et al., 2012; Karanian and Slotnick, 2017, 2018; Schacter and Slotnick, 2004). Of the few encoding studies, two have found occipital activation predicting true but not false recognition (Dennis et al., 2008; Kirchhoff et al., 2000; Pidgeon and Morcom, 2016; but see Garoff et al., 2005). Here, we not only show that visually specialized regions are engaged more when encoding true than false memories, but also characterize the visual features involved. Thus, insufficient early visual analysis at encoding leads to poor mnemonic discrimination of similar lures. This may prevent later recollection of details of the studied item that would allow people to reject the similar lures (recollection rejection; Brainerd et al., 2003). The RSA result is also consistent with the behavioral increase in false recognition for more visually confusable objects (see also Naspi et al., 2020).

We did not find any evidence here that semantic processing contributes to false memory encoding, and in FG, feature semantic representations impacted true memory encoding more strongly. Clearly, we cannot place weight on the null result, and our models did not comprehensively address all potential semantic processes but focused on concept-level processes we have shown to contribute behaviorally in this task (Naspi et al., 2020). Lateral and ventral temporal regions previously implicated in false memory encoding in verbal tasks did not show significant effects here (Dennis et al., 2007; Chadwick et al., 2016). These areas may support higher-level verbal semantics linking studied items to lures. Nonetheless, both in the current task and following deep semantic judgments at encoding (Naspi et al., 2020), concept confusability reduced lure false recognition relative to novel objects as well as true recognition. An intriguing possibility is that the semantic processes reducing lure false recognition operate at retrieval rather than at encoding. This hypothesis will be tested using RSA analysis of retrieval phase brain activity in this task.

In conclusion, we have revealed some of the visual and semantic representations that allow people to form memories of specific objects and later reject similar novel objects. This is the first – to our knowledge – preregistered study of neural representations in memory encoding, and the first probe of representations predicting false recognition. Using previously validated representational models, we were able to disentangle low-level image properties from semantic feature processing. The data provide novel support for theoretical assumptions implicating visual detail in specific memory encoding, but suggest that semantic information may contribute to specific as well as gist memory. Our approach offers a path by which future studies can evaluate the respective roles of encoding and retrieval representations in true and false memory.

## Acknowledgments

P.H was supported by a BBSRC New investigator grant (BB/T004444/1) Author contributions: L.N., P.H., B.D., and A.M. designed research; L.N. performed research; L.N. analyzed data; L.N., P.H., B.D., and A.M. wrote the paper

